# Characterization of the consensus mucosal microbiome of colorectal cancer

**DOI:** 10.1101/2021.06.02.446807

**Authors:** Lan Zhao, Susan M. Grimes, Stephanie U. Greer, Matthew Kubit, HoJoon Lee, Lincoln D. Nadauld, Hanlee P. Ji

**Author notes:** **Corresponding author** Hanlee P. Ji, CCSR 1115, 269 Campus Drive, Stanford, CA-94305, USA.

## Abstract

Recent evidence suggests that dysbiosis, an imbalance of microbiota, is associated with increased risk of colorectal cancer. Diverse microbial organisms are physically associated with the cells found in tumor biopsies. Characterizing this mucosa-associated microbiome through genome sequencing has advantages compared to culture-based profiling. However, there are notable challenges in accurately characterizing the features of tumor microbiomes with methods like transcriptome sequencing. Most sequence reads originate from the host. Moreover, there is a high likelihood of bacterial contaminants being introduced. Another major challenge is the microbiome diversity among different studies. Colorectal tumors demonstrate a significant extent of microbiome variation among individuals from different geographic and ethnic origins. To address these challenges, we identified a consensus microbiome for colorectal cancer through analyzing 924 tumors from eight independent RNA-Seq data sets. A standardized meta-transcriptomic analysis pipeline was established and applied to the complete CRC cohort. Common contaminants were filtered out. Our study involved taxonomic investigation of non-human sequences, linked microbial signatures to phenotypes and the association of microbiome with tumor microenvironment components. Microbiome profiles across different CRC cohorts were compared, and recurrently altered microbial shifts specific to CRC were determined. We identified cancer-specific set of 114 microbial species associated with tumors that were found among all investigated studies. Validating our approach, we found that *Fusobacterium nucleatum* was one of the most enriched bacterial species in CRC. Firmicutes, Bacteroidetes, Proteobacteria, and Actinobacteria were among the four most abundant phyla for CRC microbiome. Signficant associations between the consensus species and specific immune cell types were noted. Our results are available as a web data resource for other researchers to explore (https://crc-microbiome.stanford.edu).

## INTRODUCTION

Tumors such as colorectal cancer have specific biological interactions with commensal microbial species. Humans coexist with a rich diversity of bacteria and viruses living within the confines of specific tissue niches. This collection of microbial organisms, referred to as the microbiome, vastly outnumber the eukaryotic cells making up our various tissues [1, 2]. The cellular interactions of specific organ tissue and the microbiome can be beneficial, neutral or pathogenic in terms of non-infectious human diseases. Beneficial microbes play critical roles in maintaining immune function, metabolic homeostasis, and overall health [3]. Neutral bacteria have no discernible consequences on the host. Pathogenic microorganisms may increase the risk and severity of conditions like inflammation [4], obesity [5], fatty liver disease [6], type 2 diabetes [7] and carcinogenesis [8]. An indicator of a microbial influence in disease pathogenesis, dysbiosis is an imbalanced state of the naturally-occurring microbiota where specific pathogenic microbes overgrow other components. This phenomenon leads to a fundamental shift in the contents of the microbiome. This imbalance has the potential to lead to cancer [4]. The microbiome properties of colorectal cancer **(CRC)** have been of interest given that the colon and the rectum have the most abundant and diverse microbiome for any human organ. Many studies seek to identify specific microbiome properties that are indicators of dysbiosis and influence colorectal cancer development, phenotype and clinical outcomes.

There are two specific environmental niches for the analysis of the colorectal cancer microbiome. Generally, the largest and most diverse niche involves the microbial and viral contents of the fecal material within the colon. The smaller niche, a subset of the fecal material, involves those microbes that are directly associated with the colorectal tumor and the surrounding normal colon mucosa. This mucosa-associated microbiome has an important role in colorectal cancer biology given its direct contact to the colon epithelial tumor cells and its interactions with the local tumor microenvironment **(TME)**. Because of its direct contact with the colon cellular microenvironment, this microbiome niche is carried over after a biopsy or surgical resection of a tumor. Thus, the tumor extracted DNA or RNA reflect the microbial contents adjacent to and intermingled with the colon mucosa.

For genomic microbial characterization of CRCs, next generation sequencing **(NGS)** methods like RNA-Seq have been used for determining the microbiomes of specific tissues. For example, Simon *et al* investigated more than 17,000 samples from publicly available human RNA-Seq data and found that a significant proportion of unmapped reads were of microbial origin [9]. Sequencing the 16S rRNA gene is another common method for determining microbiome characteristics. The 16S gene contains nine hypervariable regions (V1-V9) that provide a sequence barcode for identifying microbial species and conducting phylogenetic analysis [10]. Depending on the sequencing approach, microbial abundance estimation is represented in operational taxonomic units **(OTUs)** or amplicon sequence variants **(ASVs)**, which are usually mapped to the genus or species level [11]. Each molecular dataset captures different aspects about the patient’s microbiota; comparative analysis of data from these two methods may provide insights not possible through a single data type alone.

For either the fecal or mucosal-associated microbiome, there is substantive evidence that dysbiosis is associated with the development and progression of CRC [10, 12, 13]. Studies have focused on either studying (1) the fecal contents from CRC patients or (2) direct analysis of CRC tumors with the microbiome that is in direct contact with the tumor. Citing a study from the former, Sobhani *et al* [12] performed one of the first studies to identify cancer-related dysbiosis in CRC from the analysis of fecal material from patients. They found that an elevated representation of the *Bacteroides/Prevotella* genus was present among the majority of the CRCs they investigated. Using a similar approach, Yu *et al* [13] did a metagenomic profiling of CRC samples and showed that four microbial species, including *Parvimonas micra, Solobacterium moorei, Fusobacterium nucleatum* and *Peptostreptococcus stomatis* were enriched in individuals with CRC compared to normal controls. These studies were limited to fecal samples which represent a distinct niche from CRC tissue samples.

The direct sequencing analysis of CRC tumors and mucosa provides insight into the microbiota that are directly associated with the TME. Given their proximity to the cellular milieu of the tumor, these microbes may play a potential role in the physiopathology of CRC [14]. Citing the most widely validated example of mucosa-proximal microbiome of CRC, many studies have demonstrated an enrichment of *Fusobacterium nucleatum*, which we will refer to as *F. nucleatum* for short, in CRC tumors. The initial discoveries were based on identifying microbial sequence reads from genomics studies of CRCs [15]. Some studies have shown that *F. nucleatum* is associated with higher stage CRC and a lower density of T-cells in the CRC TME. Some of these observations have been born out experimentally [16]. For example, this bacteria activates the WNT signaling pathway in CRC cells and inhibits T-cell-mediated immune responses against tumors [17].

Obtaining a high-quality analysis of cancer microbiomes has a number of significant challenges. In the case of the mucosa-associated microbiome, samples are exposed to contamination across multiple steps as a clinical biopsy is acquired, processed and sequenced. This includes the presence of microbial DNA among the molecular biology reagents used sequencing and genetic characterization. Complicating any analysis, the use of stringent quality controls has been inconsistent for cancer-based genomic studies of the microbiome [18]. These issues can dramatically skew microbiome results. Different sequencing methods such as 16S and RNA-Seq reveal different microbiome features. As an added challenge, microbiomes vary among individuals living in different geographic regions and ethnic backgrounds. This fundamental variation among individual microbiomes makes it more difficult to identify common microbial species that may have a universal role in colorectal cancer tumorigenesis.

Addressing these challenges, we analyzed the colorectal cancer microbiome composition and potential function in modulating cancer and the immune system. More importantly, we sought to identify consensus mucosal species that were consistently observed across multiple independent CRC cohorts. We utilized different RNA-Seq datasets including the Cancer Genome Atlas Colon Adenocarcinoma **(TCGA-COAD)** and the Gene Expression Omnibus **(GEO)** database. In total, 924 CRCs were included in this study to investigate the different microbiome profiles across different studies. With this large number of samples, we conducted a rigorous quality control to eliminate potential contaminants and reduce the effect of batch bias. To evaluate the quality of our mucosa-associated RNA-Seq data in evaluating microbiomes, we compared these results to a 16S analysis for a subset of overlapping samples. Finally, we derived a consensus microbiome composition across different CRC cohorts, determined dysbiosis features when examining normal tumor pairs and investigated several microbial species’ association with CRC’s cellular and clinical metrics. To facilitate the sharing of this consensus microbiome, our results are available and can be queried through a web data resource (https://crc-microbiome.stanford.edu).

## METHODS AND MATERIALS

### Colorectal tumor RNA-Seq data

Seven whole-transcriptome sequencing CRC datasets were downloaded either from NCI’s Genomic Data Commons (**GDC**) or the Sequence Read Archive **(SRA) (Table 1)**. In addition, we had an internal data set from an independent set of CRCs that we refer to as **IMS3**. All participants signed a written informed consent as part of a study protocol approved by Stanford University. Tumor tissues were collected and preserved on formalin-fixed paraffin-embedded **(FFPE)** slides. All tumor samples were determined to have greater than 60% cellularity in pathology review.

**TABLE 1.** RNA-Seq datasets included in the study.

| Study | Sequencing Platform | Sample Origins | Sample Type | Sample Size | Species Identified |
| --- | --- | --- | --- | --- | --- |
| IMS3 | Illumina | Western United States | **Tumor (N=924)** | 162 | 731 |
| TCGA | Illumina | Multiple countries |  | 564 | 4187 |
| GSE107422 | Illumina | South Korea |  | 109 | 744 |
| GSE146889 | Illumina | Midwest United States |  | 42 | 1293 |
| GSE50760 | Illumina | South Korea |  | 18 | 951 |
| GSE95132 | Illumina | Eastern United States |  | 10 | 1321 |
| GSE104836 | Illumina | China |  | 10 | 1729 |
| GSE137327 | BGISEQ | Eastern United States |  | 9 | 3378 |
| IMS3 | Illumina | Western United States | **Matched normal colon (N=298)** | 162 | 635 |
| TCGA | Illumina | Multiple countries |  | 51 | 3763 |
| GSE146889 | Illumina | Midwest United States |  | 38 | 1412 |
| GSE50760 | Illumina | South Korea |  | 18 | 883 |
| GSE95132 | Illumina | Eastern United States |  | 10 | 1395 |
| GSE104836 | Illumina | China |  | 10 | 1673 |
| GSE137327 | BGISEQ | Eastern United States |  | 9 | 3305 |

### DNA and RNA sequencing

Tumor tissues were recovered and processed for nucleic acid. RNA was extracted from Maxwell 16 LEV RNA FFPE Purification Kit (Promega, Wisconsin, USA) following the manufacturer’s instructions. RNA-Seq libraries were prepared using KAPA RNA HyperPrep Kit with RiboErase (HMR) (Roche, California, USA) by 8 cycles of PCR. The enriched libraries were quantified by qPCR using Kapa Library Quantification kit (Roche, California, USA), and subjected to Illumina MiSeq sequencing (100 bp paired-end reads).

DNA was extracted using the Promega AS1030 Maxwell 16 Tissue DNA Purification Kit (Promega, Wisconsin, USA) following the manufacturer’s protocols. The concentration of DNA was quantified with the Qubit system (Thermo-Fisher Scientific, Massachusetts, USA), and DNA integrity was evaluated using LabChip GX (PerkinElmer, Waltham, Massachusetts, USA). Five hundred ng DNA from each sample was sheared using a Covaris E220 sonicator (Covaris, Massachusetts, USA) (microTUBES AFA fibre, 10% duty cycle, 200 cbp, intensity 5, and time 55 s), and purified by a 0.8X AMPure XP (Beckman-Coulter, California, USA) bead cleanup. The hypervariable regions (V3-4) of the 16S rRNA gene from each sample were amplified using Forward primer (5’-TCG TCG GCA GCG TCA GAT GTG TAT AAG AGA CAG CCT ACG GGN GGC WGC AG-3’) and Reverse primer (5’-GTC TCG TGG GCT CGG AGA TGT GTA TAA GAG ACA GGA CTA CHV GGG TAT CTA ATC C-3’) with Illumina sequencing adaptors (Illumina, California, USA). The purified PCR products were then subjected to a multiplexing process using Nextera XT Index kit (Illumina, California, USA) in 50 μL reactions. After PCR product cleanup, two batches of libraries were quantified and sequenced using an Illumina MiSeq platform.

### Sequence data processing for microbiome characterization

Raw RNA-Seq data were preprocessed to remove adapter sequences and low-quality bases with Cutadapt (v2.4) [19]. Trimmed data were then mapped to the human genome (GRCh38) using STAR (v2.5) [20]. Uniquely mapped reads were used for subsequent immune cell infiltration analysis. Quality controlled (by fastp software) unmapped reads were used as microbial reads for taxonomic assignment for each OTU.

Reverse reads from 16S amplicon sequencing were removed from the analysis due to low sequence quality. The sequence was processed with DADA2 using maxN = 0, maxEE = 2, truncQ = 2 parameters to do reads filtering and quality checks [21]. Reads that passed the quality control were used for taxonomy classification. ASV values were determined for each sample.

### Taxonomic microbiome classification

Kraken2 was used as the meta-transcriptome classification tool in our study. It relies on exact k-mer matches to assign microbial sequences to specific taxonomic labels [22]. The unmapped reads from RNA-Seq were queried in a Kraken2 database we created on our server (September 18, 2019), which contains taxonomic information (obtained from NCBI Taxonomy database), complete genomes in Refseq for the bacterial, archaeal, viral, plasmid, and eukaryotic organisms. Taxonomy classification results were posted to the CRC consensus microbiome website (https://crc-microbiome.stanford.edu).

### Gene expression quantification and immune cell infiltration analysis

Gene counts table generated from RNA-Seq mapped reads were normalized using TMM (weighted trimmed mean of M-values) with the EdgeR package and converted to cpm and log2 transformed [23]. A filtering process was also performed to exclude genes without at least 1 cpm in 20% of the samples. We used the program Xcell to estimate 64 tumor infiltrating immune and stromal cell types, together with immune, stroma, and tumor microenvironment **(TME)** scores for each tumor’s or normal colon’s RNA-Seq data (xcell.ucsf.edu) [24]. Multiple testing correction was applied using the p.adjust() function available in R, with the method set as “FDR”. Kruskal-Wallis rank sum test was used to determine differential immune cell infiltration among patient groups using a threshold of multiple testing corrected P < 0.05.

### Differential microbiome analysis

Microbial differential analysis was performed using DEseq2 and Phyloseq [25]. Statistical tests such as the Chi-Squared test and the Wilcoxon rank-sum test were performed to examine the patient grouping information with various clinical variables. Multiple testing correction was applied as previously described. Results were considered significant if the adjusted p-value was less than 0.05.

### The CRC Microbiome Explorer website

We developed a web-based data resource for our study (https://crc-microbiome.stanford.edu). The microbial abundance data was uploaded to a MySQL (v5.5.62) relational database from kraken2 output converted to mpa format. The database server has 32GB RAM and 16 processors running Ubuntu (v16.04). The web application was written using Ruby on Rails (v5.1.7 with ruby v2.4.2), a framework well suited for use with a backend relational database. The application server uses Ubuntu (v16.04). The application was deployed using Passenger and Apache2. The user interface utilizes Bootstrap (v3.4.1) for responsive sizing to different format clients and browsers. Jquery dataTables provide standard formatting, search and filtering capability for query tables, and Highcharts is used to format and display plots. All queries and plots are produced dynamically from the underlying database tables based on user query parameters.

## RESULTS

### CRC microbiome composition estimation from unmapped RNA-Seq

Overall, we analyzed eight primary CRC transcriptomic datasets [26-32] from a variety of sources that included the Cancer Genome Atlas **(TCGA)**, the NIH Gene Expression Omnibus **(GEO)**, the NIH’s Short Read Archive **(SRA)** and an independent dataset **(IMS3)**. The CRC RNA-Seq studies included the TCGA COAD data set which had the largest number of samples (n=564). In addition, GEO had six different data sets with the highest number of CRCs coming from GSE107422 (n=109) and the smallest set being GSE137327 (n=9) **(Table 1)**. The IMS3 data set contained 162 tumor and matched normal tissues. The total number of CRC samples were 924. An additional 298 matching normal colon samples were available for assessing their microbiome characteristics. Except for GSE137327 which used the BGI sequencing technology, all of the samples were sequenced with Illumina.

To process these CRC RNA-Seq cohorts, we removed human genome sequences, low quality reads and adapter sequences. Subsequently, we used the high quality microbial (unmapped) reads from a given CRC sample for taxonomy classification with Kraken2 **(Figure 1)**. We also conducted downstream processing and leveraged an updated database that includes NCBI’s RefSeq sequence data for human, bacteria, and viruses **(Methods)**. Across this extended tumor cohort, we observed that an average of 83% of reads were uniquely mapped to the human genome per sample. Quality controlled, unmapped RNA-Seq reads averaged 4% per sample. The percentage of unmapped reads for each dataset varies from 0.05% (TCGA) to 19.86% (GSE107422) **(Supplemental Figure 1)**. Variations in the raw sequence data, unmapped reads and unmapped ratios were observed from each dataset. For example, the TCGA and GSE146889 cohorts had the largest number of total and unmapped sequences per a sample. The GSE107422 as well as GSE104836 samples set had the highest percentages of unmapped reads **(Supplemental Figure 1)**. Thus, we normalized microbial abundances to the median sequencing depth within each cohort. Our results are available for exploring and download at the following URL: https://crc-microbiome.stanford.edu.

**Figure 1.**
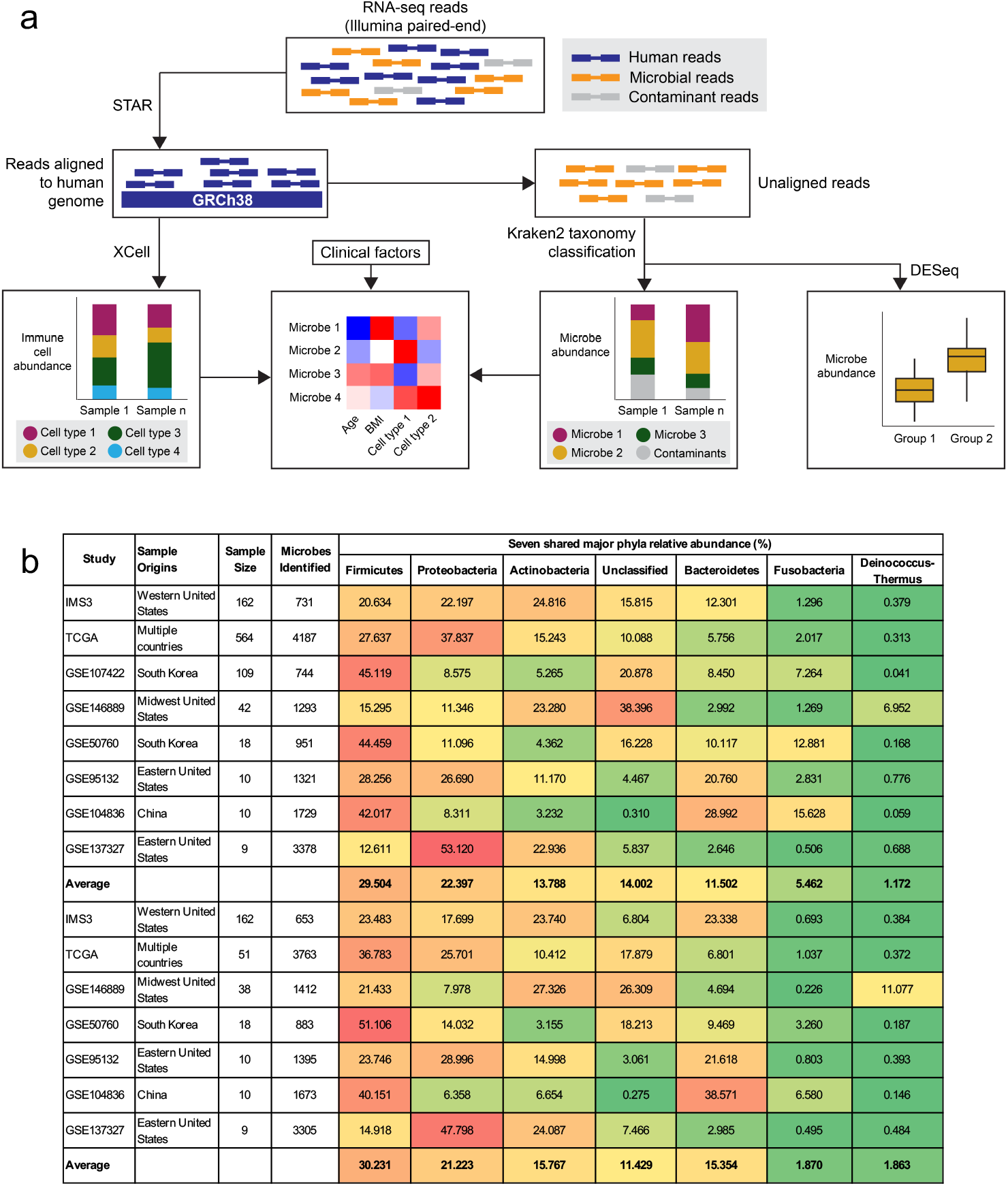
Pipeline for microbiome analysis using RNA-Seq data and analysis. (a) RNA-Seq data were processed and mapped to the Human genome. Mapped data was employed to do immune cell infiltration profiling. Unmapped reads were quality controlled and considered as microbial sequences, and used as the input to do taxonomy classification. The downstream steps included in the pipeline are microbial abundance, differential analysis and microbe-trait correlation analysis. (b) A heatmap representing the phyla determined among the eight different data sets. The phyla of the colorectal tumor and normal microbiome’s representation is shown via a relative abundance percentage each phyla across each study. Red is indicative of a higher fraction and green indicates a lower fraction.

A portion of sequencing reads may originate from contaminating microbial DNA that are found in the general environment or contaminants from the sequencing assay. This includes microbes that contaminated the sample as a result of clinical processing, were present in the sequencing reagents or grow in the fluidic systems of sequencers. To minimize the bias introduced by unwanted information, we conducted a microbial filtering process. Known microbial contaminants were filtered out **(Supplementary Table 1)**. This list was compiled by Eisenhofer *et al*. based on a series of negative controls across multiple studies. This list of contaminants was part of their ‘RIDE’ minimum standards criteria which addresses many of the potential sources of artifacts in genomic-based microbiome characterization [33].

We identified the highest represented phyla from each cohort and made comparisons of relative percentage abundance **(Table 1, Figure 2a-g)**. The shared seven phyla were displayed in the heatmap **(Figure 2h)**. Firmicutes, Proteobacteria, Bacteroidetes, and Actinobacteria were the four top ranked bacterial phyla identified from various CRC cohorts. The average relative abundance of Firmicutes (over 29.5%), had the highest average abundance across the entire cohort. This species was followed by Proteobacteria (22.4%), Actinobacteria (13.8%), and Bacteroidetes (11.5%).

**Figure 2.**
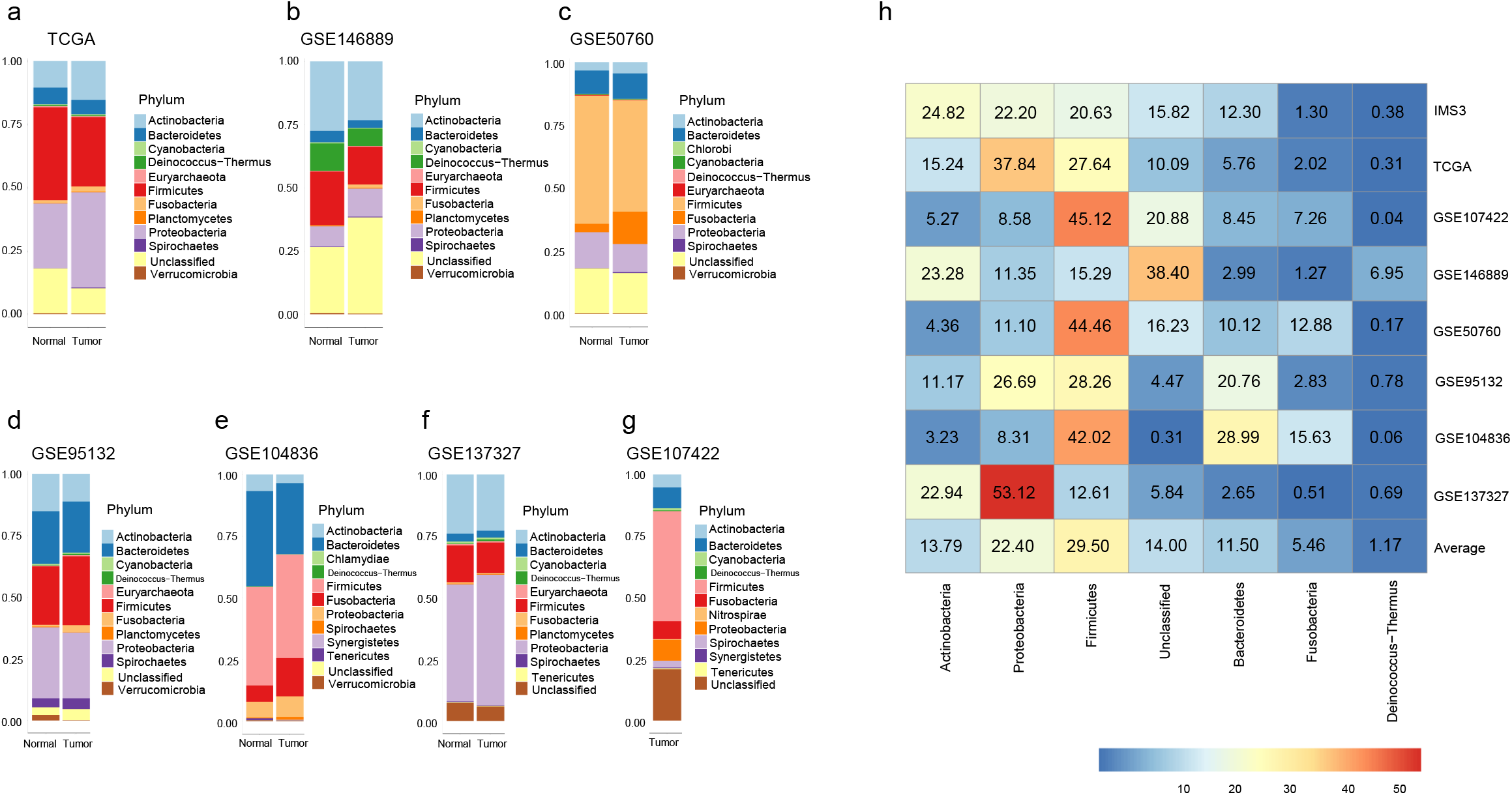
CRC microbial composition varies in different populations. The top 12 most enriched phyla identified from each cohort: TCGA (a), GSE146889 (b), GSE50760 (c), GSE95132 (d), GSE104836 (e), GSE137327 (f), and GSE107422 (g). The shared most abundant 7 phyla relative abundance plot (h). In the heatmap, columns correspond to microbes, and rows to different dataset. Relative abundances were represented by different colors, red means higher values, and green, lower ones.

Variations of bacterial community composition were observed at the phylum level, such as Proteobacteria accounts for more than half of the major phyla abundance in GSE137327, whereas this species only accounted for less than 10% in GSE104836 **(Figure 2h)**. Other noticeable phyla include Fusobacteria and Deinococcus-Thermus (5.46% and 1.17% relatively). These phyla accounted for a small proportion of the total percentages of relative abundance, respectively **(Figure 2h)**. Overall, the bacterial community composition variations were observed at the genus level **(Supplemental Figure 2)**.

### Comparing RNA-Seq versus 16S for identifying and characterizing CRC microbiome

One CRC cohort had overlapping RNA-Seq and 16S data (IMS3, n=162). We used this data for a comparison study between the two sequencing methods. The processing and analysis of microbial reads derived from RNA-seq data were described above. The raw 16S sequencing data was processed using DADA2 and phyloseq pipelines. Adapters, low quality bases and amplification primers were filtered out. Approximately 95% of 16S rRNA sequences passed our quality control measures, bringing in an average of 47,000 reads per sample for taxonomy assignments using Silva v132 annotation. DADA2 detected 531 ASVs, after removal of ASVs that were not present in at least one read count in 1% of the samples **(Supplemental Table 2)**. Fifteen and 172 bacterial taxons were observed from ASV on the phylum- and genus-level, respectively.

We compared the RNA-Seq and 16S methods as a way of evaluating the accuracy of RNA-seq-based microbiome phylum and genus level characterization. There were 12 common phyla identified from these two platforms **(Supplemental Figure 3a-b)**. Actinobacteria, Proteobacteria, Firmicutes, and Bacteroidetes were the four most prevalent **(Supplemental Figure 3a)** and abundant phyla **(Supplemental Figure 3b)**. High Pearson correlation coefficients were observed for phylum-level prevalence (0.977) and abundances (0.962) between 16S and RNA-Seq data **(Supplemental Figure 4a-b)**. The prevalence and relative abundance of Bacteroides, Nonanavirus, Actinoplanes, Bacillus, Actinetobacter, and Streptomyces were much higher than the remaining genera identified from RNA-Seq data **(Supplemental Figure 3d-e)**.

We determined that the microbial diversity via a Shannon index estimate from RNA-Seq data was significantly higher compared to the results from the 16S data at the phylum/genus levels **(Supplemental Figure 3c,h)**. Statistical significance was demonstrated using pairwise wilcox test (p<2e-16). A total of 89 overlapped genera were evident when comparing these two different methods **(Supplemental Table 3-4)**. Bacteroides and Faecalibacterium were the two most enriched genera identified both from 16S and RNA-Seq data **(Supplemental Figure 3f-g)**. The Pearson correlation coefficients between these two sequencing methods at the genus-level were 0.583 for prevalence, and 0.807 for abundances **(Supplemental Figure 4c-d)**. The differences between the two were largely due to the viral genome species that were present in the RNA-Seq data. Overall, these results not only suggest that RNA-Seq analysis of CRC accurately determines the microbiome features but also provide much more species information than 16S data. In addition, our results showed that 16S data had limited resolution beyond the genus-level and a more restricted degree of microbial characterization. Thus, we opted to focus on using the RNA-Seq data for the remainder of the study.

### The CRC consensus mucosa-associated microbiome

From the 924 CRCs and the tumor RNA-Seq data, the high-quality unmapped reads underwent Kraken2 processing and species classification. The range of species identified prior to consensus filtering was from 731 (IMS3) to 4187 (TCGA) **(Table 1)**. To determine the microbiome features that were generalizable across the entire cohort, we created a union matrix representing all of the samples and different species across the entire cohort. Subsequently, we applied a 1% prevalence filter, retaining only the bacterial and viral species above this frequency threshold. Among the eight studies, 126 microbial species were obtained from the 924 CRC tumors. We conducted an additional level of filtering to determine if identify any species typically associated with contaminant artifacts. There were a few species that are known environmental contaminants, such as microbe belonging to the the Cutibacterium and Methylobacterium genera. These contaminants and others were removed, which resulted in a final consensus list of 114 microbial species associated with CRCs **(Table 1, Supplemental Table 5)**. All of the species were present for all of the sample sets included in the study.

Bacteroidetes and Firmicutes species account for a significant proportion of the 114 microbial list **(Figure 3)**. From our consensus CRC microbiome, more than 33% of these species belong to the class of Clostridia **(Figure 3a)**. This class was the most frequently occurring among our cohort. Species belonging to Bacteroidetes (23.5%), Proteobacteria (16.5%), and Actinobacteria (10.4%) were the second, third, and fourth most predominant phyla among the consensus microbiome. Most members of the Clostridia have a commensal relationship with the host, and are involved in the maintenance of intestinal health [34]. Other well-characterized fecal species included *Bacteroides megaterium, Bacteroides fragilis, Escherichia coli, Bacillus cereus, Faecalibacterium prausnitzii, Bacteroides vulgatus*, and *Prevotella intermedia* and were among the most abundant species of all the CRC samples across the cohort **(Supplemental Table 5)**. Validating our analysis results, *F. nucleatum* was common among the tumors, has been previously associated with CRC and has a mechanistic contribution towards colon cancer growth.

**Figure 3.**
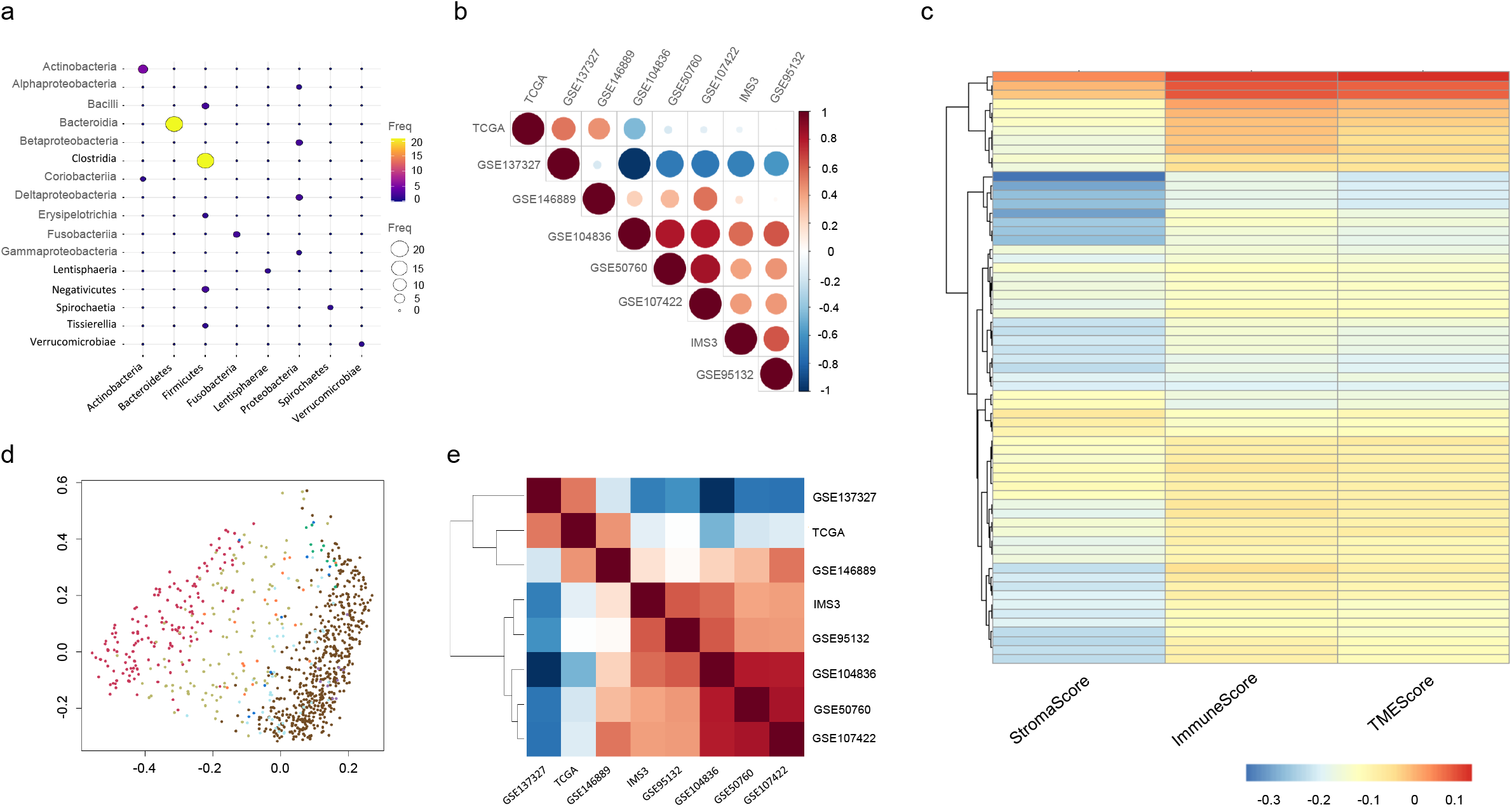
Colorectal cancer’s consensus microbial species. Balloon plot to summarize and compare the taxa distribution for the 114 species at the phylum (x-axis) and class (y-axis) levels, where the area and color of the dots were proportional to their numerical value (Freq) (a). The correlation plots (b, e) of the eight CRC cohorts. Red represent positive correlation, and blue negative correlations. (b) was a correlogram, where the area and color of the dots were proportional to their correlation coefficients. (e) was a correlation heatmap. (b) and (e) share the same color legend. (c) The PCA-based on Bray-Curtis dissimilarity was used to estimate the beta diversity of the cohorts. (d) A correlation heatmap of the 114 species with stroma, immune, and TME scores derived from the same tissue. The rows are specific microbe species found in our colorectal cancer consensus microbiome. The columns are labelled with the cell type summaries derived from the RNA-seq data.

Interestingly, a number of these microbiome species are established pathogens with the most prevalent being *Clostridium difficile* which is a major cause of infectious diarrhea. This species is also associated with inflammatory bowel disease **(IBD)** [35]. *Clostridium perfringens* is one of the most common causes of food poisoning in the United States [36]. Besides the pathogens such as *Clostridium difficile and Clostridium perfringens*, other potentially pathogens such as *Akkermansia muciniphila* is a mucin-degrading bacterium. *Pasteurella multocida* can cause a range of diseases in animals and humans, particularly for skin infections.

Other species are commensal elements of normal gut microbiota. Some of them possess probiotic properties, like *Bacteroides xylanisolvens* and *Bacteroides ovatus*. Others play important roles in other mammalian species and extrinsic metabolic processes. For example, *Lachnospiraceae bacterium* can ferment polysaccharides into short-chain fatty acids and alcohols [37]. *Bacteroides cellulosilyticus*, a strictly anaerobic cellulolytic bacterium, metabolizes cellulose to smaller molecules, and ferment various carbohydrates [38]. *Ruthenibacterium lactatiformans* is characterized by fermentative metabolism [39]. *B. megaterium* has probiotic potential [40].

### Consensus microbiome from matched normal colon tissue

From the 298 matched normal colon tissue and their RNA-Seq data, we applied the same bioinformatic process, quality control filtering and prevalence analysis with a union matrix. The range of species identified prior to consensus filtering was from 635 (IMS3) to 3763 (TCGA) **(Table 1)**. From this analysis, there were 153 species consistently found among all of the matched normal tissues **(Supplemental Table 6)**. More than half of the species were identical to the tumors’ 114 species list **(Supplemental Figure 5)**. Interestingly, the remaining consensus mucosa-associated microbiome from normal colon tissue was quite different from tumors, with a large proportion of them came from the *Proteobacteria* and *Firmicutes* phyla **(Supplemental Table 6)**. *Proteobacteria spp* occurs as a free-living species which can be identified within the colon microbiota. *Firmicutes* phyla, especially in the class of *Clostridia* were enriched in normal tissues, suggesting that *Clostridia spp*. were potentially beneficial microbes. When compared to the matched normal tissue set, 29 species were only present in tumor tissues, this include two *Fusobacteria* species and several other known pathogens (*Streptococcus spp*. and *Prevotella spp*.).

We investigated the geographic associations of the eight datasets using the abundances of the selected 114 microbial species specific to CRC tumors. The GSE137327 microbial profile was negatively associated with other cohorts, suggesting that the choice of sequencing platform may have affected the microbiome profile **(Figure 3b,e)**. This data set was generated from a different sequencing platform, the BGI-Seq system versus the remainder of the studies which were Illumina-based. Several Asian cohorts from different geographic locations were represented in this study. This included GSE107422 and GSE50760 where the tumor samples originated from South Korea. For the GSE104836 cohort, the samples originated from mainland China. The CRCs from all three of these studies were part of a distinct cluster with Asian origins **(Figure 3b, e)**.

### TME and immune cell correlations with the CRC consensus microbiome

Using the same RNA-Seq data per sample, we determined the immune cell estimations of each CRC across the cohort. Currently, one can use bulk RNA-Seq data to infer the proportions of individual cell types from tumor samples. This process is generally referred to as cell deconvolution. To conduct this study we used the program xCell to make estimates about the relative cell populations among the CRC RNA-Seq data sets [24]. This program deconvolutes gene expression data to identify the relative representation of 64 immune and stromal cell types.

After deconvoluting the gene expression data from our cohort, we determined the association of TME components that are referred to as the immune, stroma, and TME scores. For this analysis, we used the 114 tumor-specific microbial species from the CRC consensus microbiome. As an aggregate indicator of different types of cellular composition, Xcell provided each CRC with an immune score (the sum of all immune cell types), stroma score (the sum of all stroma cell types), and TME score (the sum of all immune and stromal cell types) [24]. Stroma scores were mostly negatively correlated with the 114 CRC species compared to immune and TME scores **(Figure 3c)**.

Thirty-eight microbial species were selected which have significant associations with specific types of the immune cells (Spearman correlations, FDR < 0.05) **(Figure 4a)**. Natural killer **(NK)** cells had a positive correlation with the majority of the selected microbial species. CD4 T, CD8 T, naïve/pro B, and T regulatory cells play opposite correlation patterns with NK cells **(Figure 4a)**. In other words, these cells were negatively correlated with more than half of the CRC consensus microbes. *Victivallales bacterium* (CCUG447300) was one of the species that had a significant positive correlation with NK cell’s enrichment in the CRC TME (Spearman’s rho = 0.57; p< 1e-4). This species was significantly negative correlated with the CD4 naive T cell’s abundance in the TME (Spearman’s rho = -0.45; p< 1e-4).

**Figure 4.**
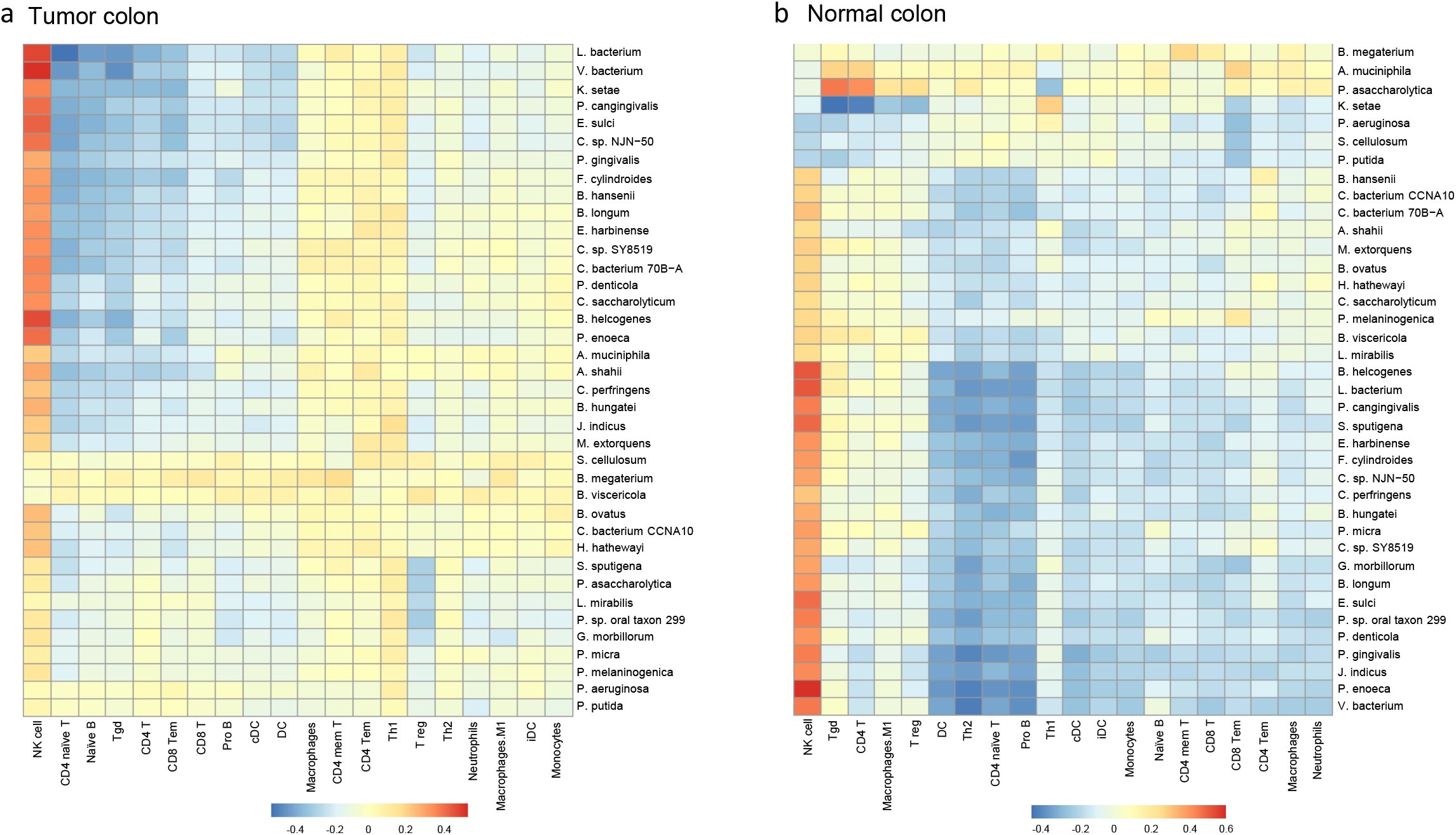
Selected microbial species correlate with immune cells. Heatmaps show the correlation patterns between microbes and immune cells in the tumor microenvironment that include (a), and normal samples (b). In the heatmaps, columns correspond to cell types, and rows to microbes. Spearman correlation values were represented by different colors, red means higher correlations, and green, lower ones.

Immune cells and microbe’s correlations were also investigated in the adjacent normal samples **(Figure 4b)**. NK cells were positively correlated with the selected thirty-eight microbial species. More negative correlations than positive correlations between immune cells and microbes can be seen from the heatmap **(Figure 4b)**. Distinct correlation patterns have been found between tumor and normal tissues. For example, macrophages and CD4 memory T cells were generally positively correlated with the selected species in tumor samples, however, the correlations changed to negative in adjacent normal tissues. Similarly, we found that T regulatory, CD4 T, T helper2, iDC, and monocytes had different correlation patterns in tumor (positive) and normal (negative) samples. A group of microbial species such as *B. helcogenes, L. bacterium, P. cangingivalis, S. sputigena*, and *E. harbinense* have shown similar trends of correlations with a subset of immune cells (DC, NK, and other innate immune cells), indicating that there were some microbe-microbe interactions between them.

### Comparison of CRC versus matched normal microbiomes

To determine the differences between matched normal and colon tissue microbiomes, we used the IMS3 and TCGA tumors. These two sample sets had sufficiently large numbers of matched normal tumor pairs to perform statistically meaningful differential analysis. We compared and identified microbial compositions between tumor and adjacent normal tissues at different taxonomic levels (phylum/genus/species).

Variations in the microbial phyla, genera, and species relative abundances were observed between tumor and normal groups, respectively **(Figure 5)**. More specifically, at the level of the phylum, increased proportion of *Fusobacteria* and virus (**Figure 5a, adjusted p < 0.01)** as well as depletion of *Bacteroidetes* (adjusted p < 0.01) **(Figure 5a)** were detected in tumors. For example, the average percentage of the viral constituents among the total tumor microbiota was 30.70% compared to 11.43% in the adjacent normal tissues. The relative abundances of Bacteroidetes (38.29% vs 22.56%) in normal tissue was detected at a higher percentage than in the tumor tissues **(Figure 5a)**. Significant differences in the abundances of three genera were observed between tumor and adjacent normal tissues **(Figure 5b)**. These genera were all under the above-mentioned phyla such as *Fusobacteria* and *Bacteroidetes*, which followed the same trend with the fold changes we observed at the phylum level.

**Figure 5.**
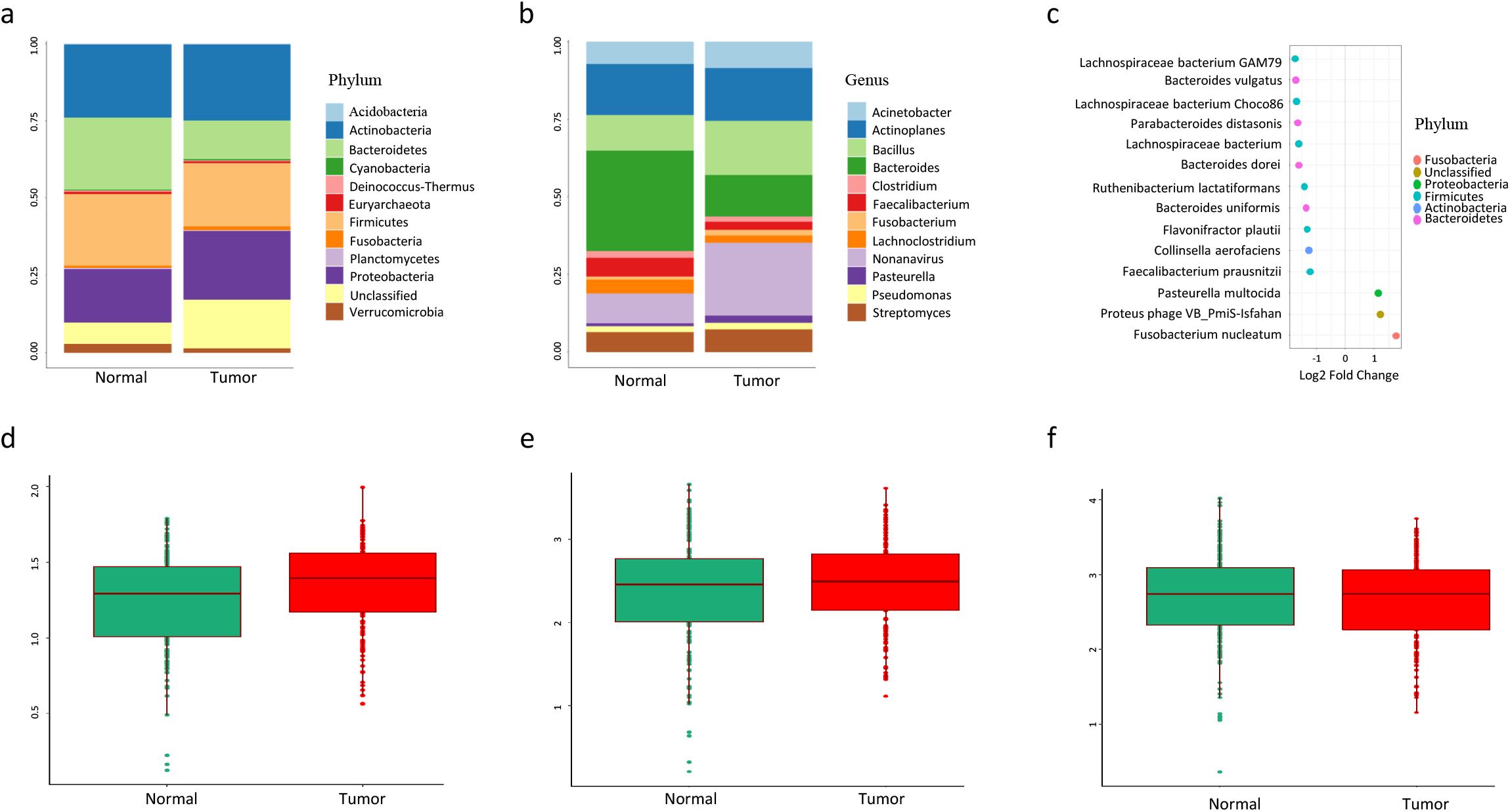
Tumor microbial composition is different from that of adjacent normal. The top 12 most abundant phyla (a) and genera (b) distribution plots for tumor and adjacent normal. Differentially enriched/depleted microbial species between tumor and adjacent normal (c), x-axis: log 2 fold changes, y-axis: microbial species names, and colors labeled their phylum levels. Comparison of Alpha diversity (Shannon index) between tumor and normal at the phylum (d), genus (e), and species (f) levels.

A total of 13 microbial species were significantly differentiated between the tumor and normal groups with an adjusted p< 0.05 **(Figure 5c, Supplemental Table 7**) in IMS3 cohort. For example, high abundance of *F. nucleatum* and *Pasteurella multocida* were identified among the tumors compared to matched normal tissue as noted by fold changes greater than 1 **(Supplemental Table 7)**. The remaining 11 species were all decreased in tumor. For instance, six members within the order *Chlostridales*, namely, three *Lachnospiraceae* and three *Ruminococcaceae* were depleted in tumors. *Lachnospiraceae* are generally beneficial microorganisms that work to fight off colon cancer by producing butyric acid [41]. *Faecalibacterium prausnitzii* from the family of *Ruminococcaceae* was notable as one of the most prominent commensal bacteria in the human gut. The remaining five species that were differentially lower in their tumor presence included *Collinsella aerofaciens*, and four members within the order *Bacteroidales* (three *Bacteroides* and one *Parabacteroides* genera). The overall diversity of the microbial community significantly decreased in tumors compared to the matched normal tissue at the species level **(Figure 5f)**. However, the microbial diversity in the tumors relative to their matched normal tissue was not significantly different at the phylum and genus levels **(Figure 5d-e)**.

From the TCGA dataset, we obtained 129 differential enriched/depleted microbial species with adjusted-p less than 0.05 in CRC **(Supplemental Table 8)**. Among them, 7 species (*F. nucleatum, Faecalibacterium prausnitzii, Fusobacterium plautii, Ruthenibacterium lactatiformans, Lachnospiraceae bacterium, Lachnospiraceae bacterium Choco86*, and *Bacteroids vulgatus*) overlapped with the 13 species we identified from the IMS3 cohort. *F. nucleatum* was found to be enriched in the tumor tissues in both the TCGA and IMS3 datasets, whereas the remaining 6 species were all depleted in tumors.

### A web-based CRC Microbiome Explorer Interface

To enable access to the study’s results, we created an interactive database entitled the CRC Microbiome Explorer (https://crc-microbiome.stanford.edu/). The CRC Microbiome Explorer enables the user to query the database by “Study” or “Patient”. The “Study” query displays an overview bar plot of the top 12 microbial phyla across all normal (if available) and tumor samples in the queried study. Additionally, users can view a bar plot that displays the microbiome composition of each individual patient in the study. Alternatively, the user can select a specific patient to submit for a “Patient” query that generates a Sankey plot displaying the top 10 microbial genii in each patient sample. The patient-level microbial abundance data is also available as a searchable and sortable table. Kraken2 output files from each study are available for download as tar archived files from the “Data Download” tab.

## DISCUSSION

The human microbiome is associated with human health, and dysbiosis can lead to a variety of disease such as colon cancer [42]. The colon is the site of one of the most diverse human microbiomes [3, 43]. CRC is a heterogeneous malignancy with distinct molecular features and clinical outcomes among patients. Besides genetic alterations, the gut microbiome may play a role in CRC initiation and progression [10, 12, 44, 45]. Most studies on CRC microbiota so far are conducted on fecal samples, which are obtained through non-invasive methods and are widely available compared to tissue samples. When considering the examination of the fecal versus mucosa-associated microbiomes, the analysis of tissues is more directly related to the microbiota contributions to the cellular physiopathology of CRC [14]. Thus, studying the microbiome in direct contact with the CRC’s microenvironment is important for revealing potential interactions and relationships. Moreover, microorganisms in the gut microbiota interact which changes the representation of any given species. In addition, it is estimated that more than 60–80% of the microbes are nearly impossible to culture using conventional microbiology techniques [14]. Thus, culture-independent analysis using high-throughput sequencing provides an opportunity to identify species that otherwise would be missed. Overall, we conducted this NGS-based microbial study including a series of different CRC tissue cohorts to identify tumor specific microbial profiles for future clinical use. We also investigated infiltrated immune and stroma components in the TME as the phenotype of interest to link them with the marker microbial species we identified. Our results are available at a genomic web resource for the CRC consensus microbiome (https://crc-microbiome.stanford.edu).

Tumor-promoting effects of the microbiome in CRC occurs through a dysbiosis mechanism, rather than by infections with specific pathogens [8]. This is different from the role of *Helicobacter pylori* in the pathogenesis of gastric carcinoma [46], where bacteria is widely recognized as a microbial carcinogen and the most important known risk factor for GC. Through our analysis, we found that CRC patients are characterized by the enrichment of a set of microbes which can have pathogenic effects in some circumstances as well as depletion of health-related microorganisms. For example, we identified 13 differentially enriched/depleted microbes using 162 paired tumor and normal tissues samples. Among them, *F. nucleatum* has been detected as a predominant species in tumors, which match well with previous studies. Several members of Clostridia possess the properties of fermenting diverse plant polysaccharides, which are beneficial to human health, were found to be depleted in CRC tissues.

We defined a consensus CRC microbiota by searching the most prevalent microbial species across several different cohorts, which can be a valuable resource for future studies. Importantly, this set of microbes are present regardless of the patients’ origins over a diverse range of geographic locations and ethnicities. This consensus represents species that may interact with the cellular tumor microenvironment of CRC. As an additional evaluation of the quality of this consensus microbiome, we determined if these species had been previously reported in in the literature as component species of the normal colon microbiome.

The connection between microbiome and CRC is likely to be bidirectional: microbiome changes may happen because of CRC development but may also contribute to CRC progression [47]. Integration information across datasets provided key insights into the gut microbiota of CRC patients. Humans have a long history of using microbes in daily life, research, and other beneficial capacities. For example, yeasts have been used for thousands of years in the production of food and beverage for human use. In scientific research, *Escherichia coli* and yeasts serve as important model organisms especially in biotechnology and molecular biology for decades. Furthermore, microbes have been used in industrial processes such as waste water treatment, and industrial chemicals/enzymes production. More recently, antimicrobial therapies have been used for patients who are carrying harmful species, and probiotic therapies for patients who are suffering from a lack of beneficial microbes. Development of novel microbiome-related diagnostic tools and therapeutic advances may become routine in the near future.

In conclusion, we identified tumor-specific bacteria patterns and signatures, which might serve as biomarkers for the prognosis of CRC. Our future works include identified prognostic microbial signatures across various cancer types, and translating the microbiome biomarkers to the clinic.

## AVAILABILITY

All of our results are available from the following URL: (https://crc-microbiome.stanford.edu). Sequence data is available at the TCGA COAD study from the NCI’s Genomic Data Commons website: https://portal.gdc.cancer.gov/projects. Additional data sets were available from the NIH’s GEO website from the following studies and their GEO identifiers (GSE107422, GSE146889, GSE50760, GSE95132, GSE104836, GSE137327). The scripts used in this study are available in an online repository (https://github.com/sgtc-stanford/crc-microbiome).

## ACKNOWLEDGEMENT

The work is supported by an award from the Fund for Innovation in Cancer Informatics (LZ, HL, HPJ) and the National Institutes of Health (2R01HG006137-04 to SUG, SMG and HPJ). Additional support came from the Clayville Foundation (SUG, SMG and HPJ).

## AUTHOR CONTRIBUTIONS

L.Z., H.L. and H.P.J. designed the study. M.K. conducted sequencing. G.S and C.S. optimized the isolation process. L.D.N. oversaw the clinical and samples resources for a the IMS samples. L.Z. developed the bioinformatic pipelines and analysis algorithms. L.Z., S.M.G., H.L. and H.P.J. analyzed the data. S.U.G. and S.M.G developed the website resource. L.Z., S.U.G. and H.P.J. wrote the manuscript.

