## Supplementary Figures for "Characterization of the consensus mucosal microbiome of colorectal cancer"

**TITLE**

**Institutions**

[1]Division of Oncology, Department of Medicine, Stanford University School of Medicine, Stanford, CA, United States

[2]Stanford Genome Technology Center, Stanford University, Palo Alto, CA, United States

[3]Intermountain Precision Genomics Program, Intermountain Healthcare, Saint George, UT, 84790, United States

**Corresponding author**

Hanlee P. Ji

CCSR 1115, 269 Campus Drive, Stanford, CA-94305, USA

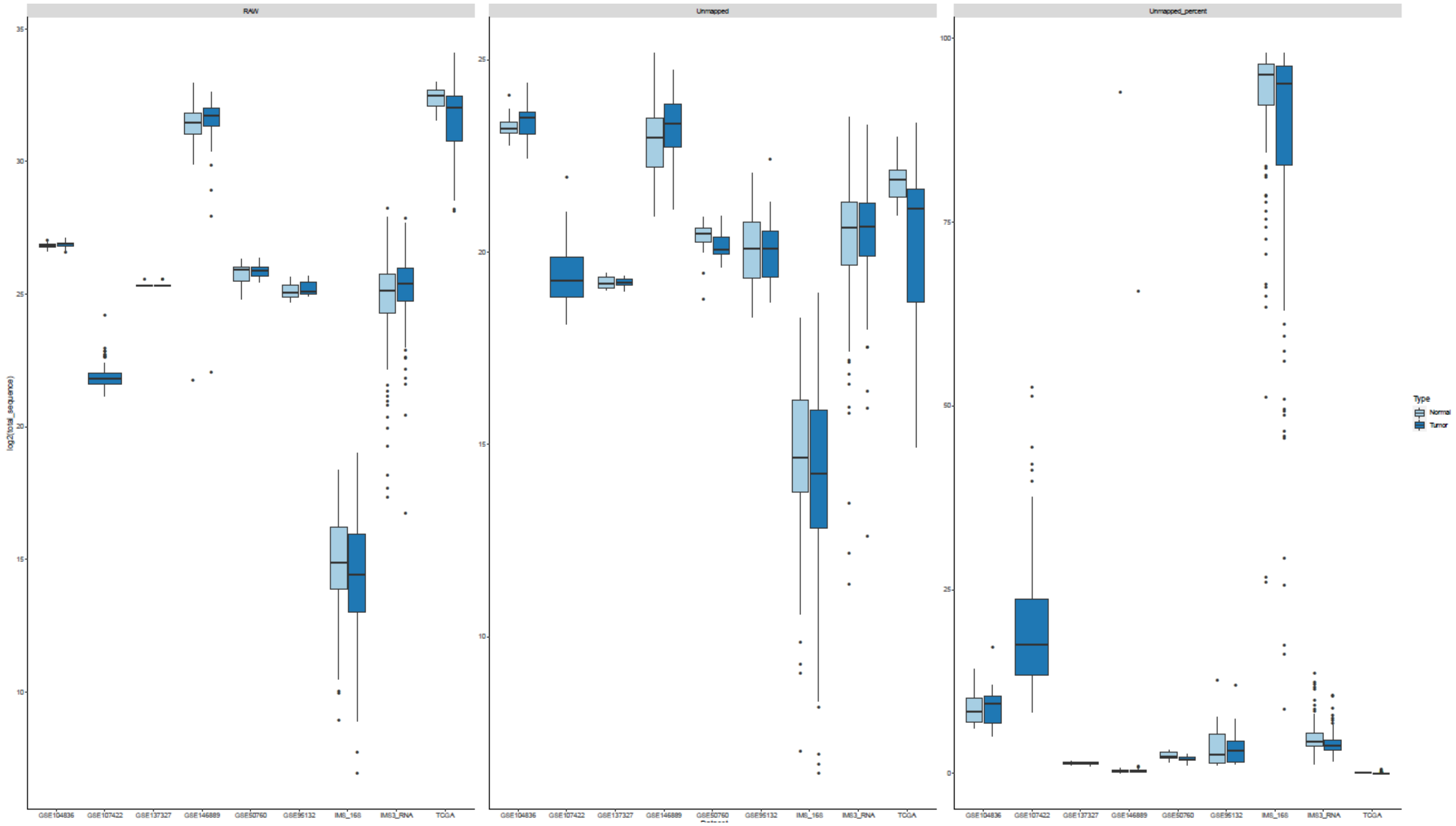


**Supplemental Figure 1. Sequence reads tracking for each dataset involved in the study.** X-axis are the names of the eight datasets. The Y-axis is for the RAW and Unmapped panels are the log2 scaled total sequences; Y-axis for the `Unmapped_percent` panel indicates the percentages of the unmapped reads. The values in the Y-axis were plotted with standard error bars for each sample set included in a dataset.

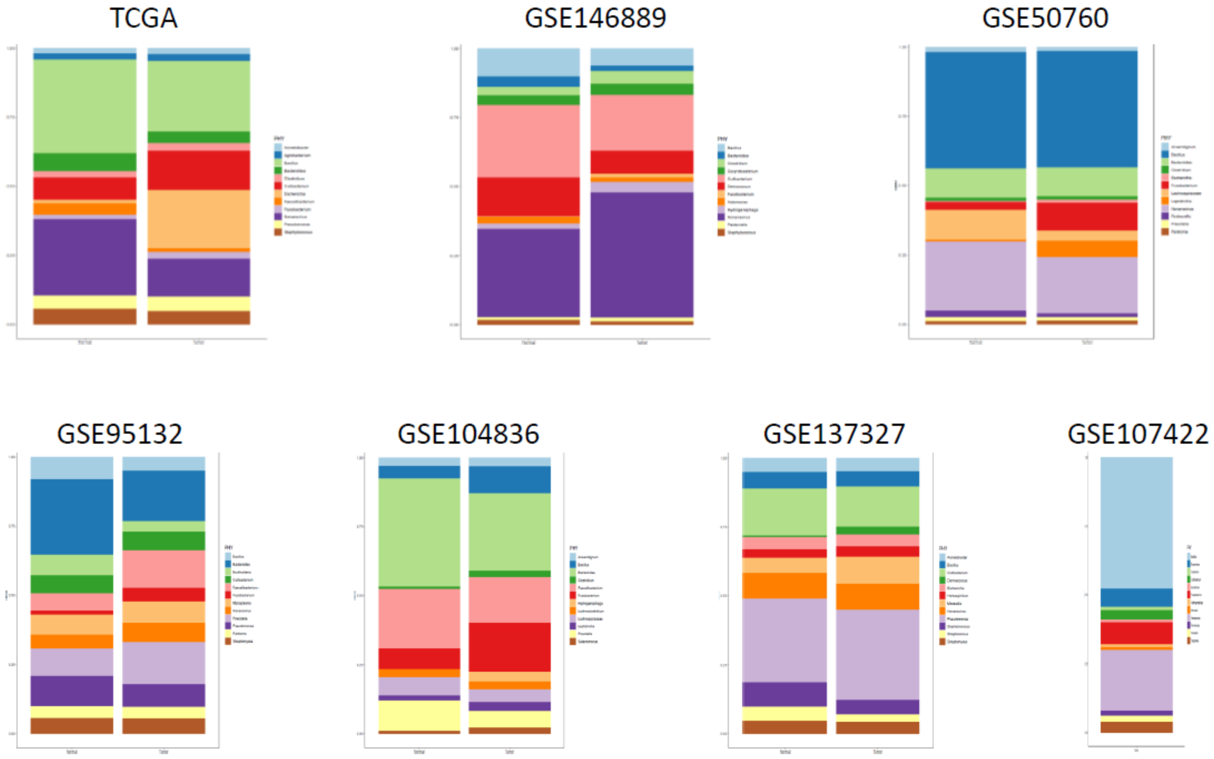


**Supplemental Figure 2. Genus-level microbial compositional changes for each dataset.** The top 12 most enriched genera identified from each cohort: TCGA, GSE146889, GSE50760, GSE95132, GSE104836, GSE137327, and GSE107422.

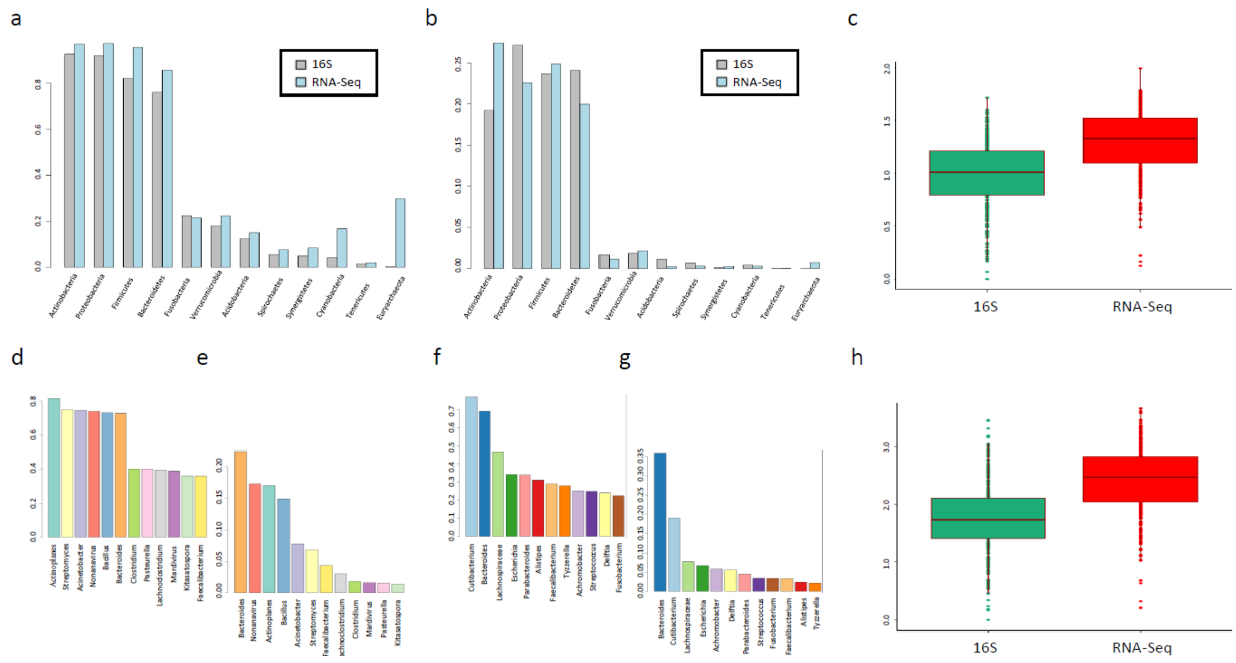


**Supplemental Figure 3. Microbial compositional and abundance comparisons between 16S and RNA-Seq at the Phylum- and Genus-level.** Prevalent (a) and abundant (b) phyla values identified from 16S and RNA-Seq platforms. Most prevalent (d-e) and abundant (f-g) phyla identified from 16S and RNA-Seq platforms. The boxplots of microbial diversity through the Shannon index comparisons at the phylum (c) and genus (h) levels.

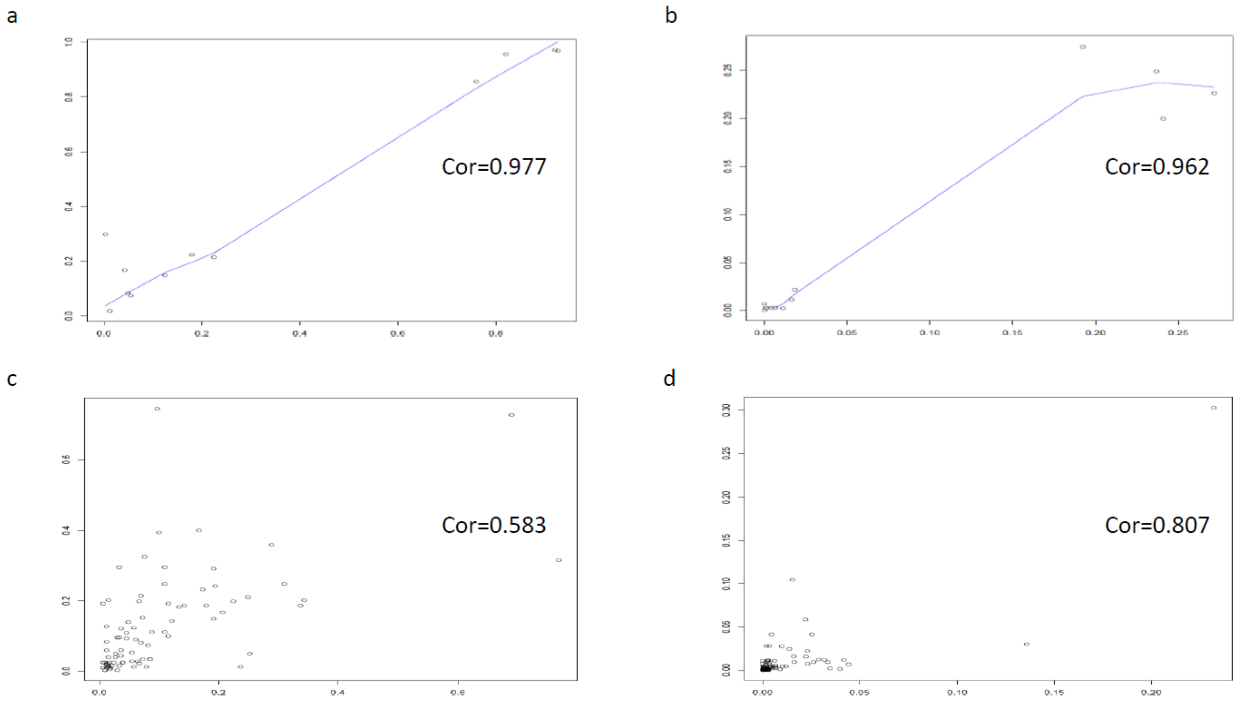


**Supplemental Figure 4. Pearson correlation coefficients between 16S and RNA-Seq at the Phylum- and Genus-level.** Phylum-level prevalence (a) and abundances (b) between 16S and RNA-Seq data; Genus-level prevalence (c) and abundances (d) between 16S and RNA-Seq data.

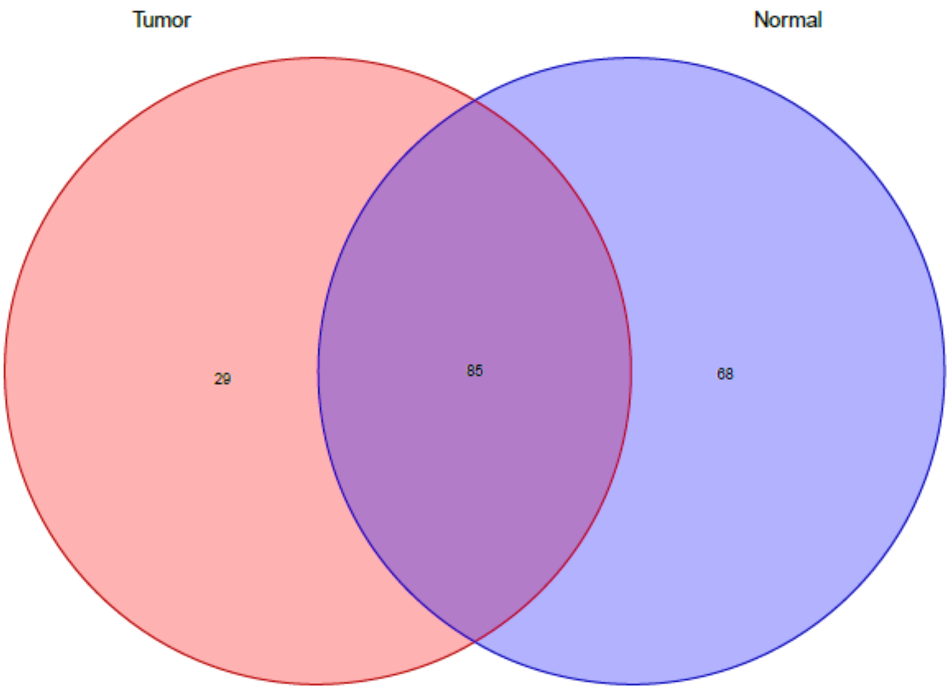


**Supplemental Figure 5. Venn diagram of the consensus microbial species between tumor and normal samples.** Venn diagram showing the number of overlapped and unique species between tumor and normal groups.
